# Comparative study of chlorophyll measurement in *Physcomitrium patens* moss using a conventional microscope adapted for combined 2D+1D imaging and spectral analysis

**DOI:** 10.64898/2026.08.27.747650

**Authors:** Pachara Thonglim, Nathan Hagen, Prathan Buranasiri, Yosuke Tamada, Rajeev Ranjan

## Abstract

Imaging spectroscopy often requires expensive and complex equipment. Here we show a simple procedure for attaching a standard miniature fiber spectrometer to a conventional microscope, allowing easy integration of 2D imaging with 1D high-resolution spectral measurements. This combination provides much of the benefit of a full imaging spectrometer without the large equipment investment, and we provide instructions for modifying microscopes to this setup and the present measurements of living cells that demonstrate their performance. Using this setup, we compare the quantitative measurement of chlorophyll concentration in *Physcomitrium patens* moss using color imaging and spectral sampling.

## 1. Introduction

Different biological tissues exhibit distinct optical absorption, scattering, and fluorescence properties, and measuring the wavelength dependence of these properties provides a way to differentiate the components of complex tissues [1, 2]. Hyperspectral imaging systems are common tools for this type of biological tissue analysis, providing spectral data at every pixel within an image [3–7]. Unfortunately, these instruments tend to be bulky and expensive, suffer from motion artifacts when measuring dynamic samples, and have low light efficiency [8]. The proposed instrument trades off the full performance of an imaging spectrometer for simplicity and ease of use, while also providing a much higher spectral resolution than is generally available among imaging spectrometers.

Although many spectroscopic techniques such as Raman microscopy provide combined spatial and spectral information, we have been unable to find research articles that explicitly address the simple and low-cost integration of two-dimensional spatial imaging with one-dimensional high-resolution spectral measurement – what we call “2D+1D measurement” within a conventional microscope. This approach offers distinct advantages: the two-dimensional image provides spatial context and enables precise localization of regions of interest within the sample, while the corresponding one-dimensional spectral data delivers detailed chemometric information from the central measurement region [9]. In many microscopic spectral analysis applications, acquiring full spectral data across the entire field of view is unnecessary; instead, high spectral resolution is typically required only in specific, targeted regions. Consequently, the proposed 2D+1D strategy represents a more efficient and focused alternative to full hyperspectral imaging, reducing data volume and computational demands while preserving essential analytical capabilities [10].

In the literature, it is common to think of standard imaging as 2D (x,y), often with color, and spectral imaging as 3D (x,y,λ), but it is easy to forget that there is a useful continuum between the two. In the discussion below, we describe a technique of sampling the spectrum at one point within the image while simultaneously collecting the full image. We informally nickname this technique as “2.1D spectral imaging” (the “point one” of “2.1” being an inversion of “one point”, which makes it easy to remember) — a naming inspired by the informal “2.5D” nomenclature widely used to describe the measurement of 3D surfaces (as opposed to 3D volumes) [11]. This approach allows a user to collect video data at the full imaging speed and quality of the original imaging system with only a modest loss in light collection. Adding a fiber spectrometer to an existing imaging system is not a new concept [12–15], but it is also not widely used. We suspect that this is partly because many do not realize just how easy it is to modify a conventional microscope for 2.1D spectral imaging. For that purpose, we describe the process in detail and give instructions for the mechanical mounting components. Another reason we suspect plays a role in why 2.1D spectral imaging is so little used is that many researchers are likely unfamiliar with just how useful a technique it is. In many of our day-to-day tasks of investigating samples, we often rely on analyzing spectra evaluated at individual locations of interest, without needing to have fully spatially resolved spectra. Using a full spectral imaging instrument for this kind of standard work is unnecessary.

In this manuscript, we provide details on how to modify a conventional microscope to enable this setup and describe the performance of the resulting system. In order to show that this modification has little effect on microscope performance, we compare measurements of a resolution test target before and after attaching the adapter. Finally, we show images and spectra of living *Physcomitrium patens* tissue, from which we quantify chlorophyll concentration using both the image and spectral data separately.

## 2. Instrument design and assembly

In order to achieve 2D+1D imaging and spectral imaging within a conventional microscope, we modified its optical layout by adding mechanical mounting components and optical elements to the side port of an inverted conventional microscope. For most conventional microscopes, there is sufficient space to add a beamsplitter between the exit port flange and the camera image plane, and this is all we need to send a portion of the light to a fiber-coupled spectrometer.

The mechanical components (see Fig. 1) are designed to allow the insertion of a beamsplitter, to move the camera to move farther away from the exit port (since the beamsplitter causes the image plane to move back by 16.4 mm), and to provide a mount point for the spectrometer’s fiber. The components were designed using computer-aided design software (SolidWorks). Three separate mechanical parts were developed: (a) a component for mounting the beamsplitter and connecting the new optical elements, (b) a component for coupling to the exit port of the microscope, and (c) a fiber-optic mount. The CAD models of the three mechanical components are shown in Fig. 1. After the CAD files were established and verified for suitability and compatibility with other components, the mechanical components were fabricated from aluminum.

**Figure 1.**
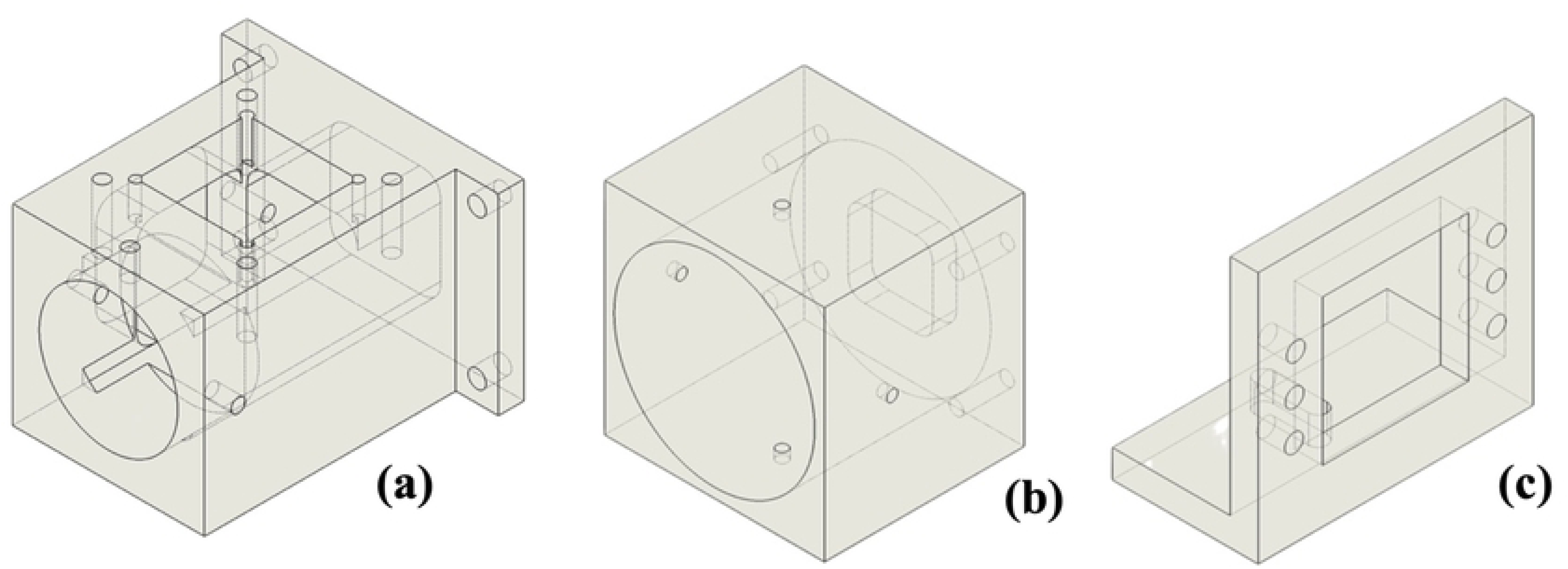
The CAD models of mechanical components of the adapter: (a) the beamsplitter mount, (b) the exit port coupler, and (c) the fiber mount coupler.

The modified microscope setup is shown in Fig. 2, where we have attached our adapter to an Olympus IMT-2 inverted microscope [16, 17]. First, we removed the MTV-3 C-mount adapter with a 0.3X reducing relay lens at the microscope’s exit port, which creates space for the beam splitter and futher extends the focal distance. The original microscope uses the 0.3X reducing relay lens to shrink the image from the much larger film standard size (36×24 mm) approximately to a 1-inch digital format (∼12×8 mm).

**Figure 2.**
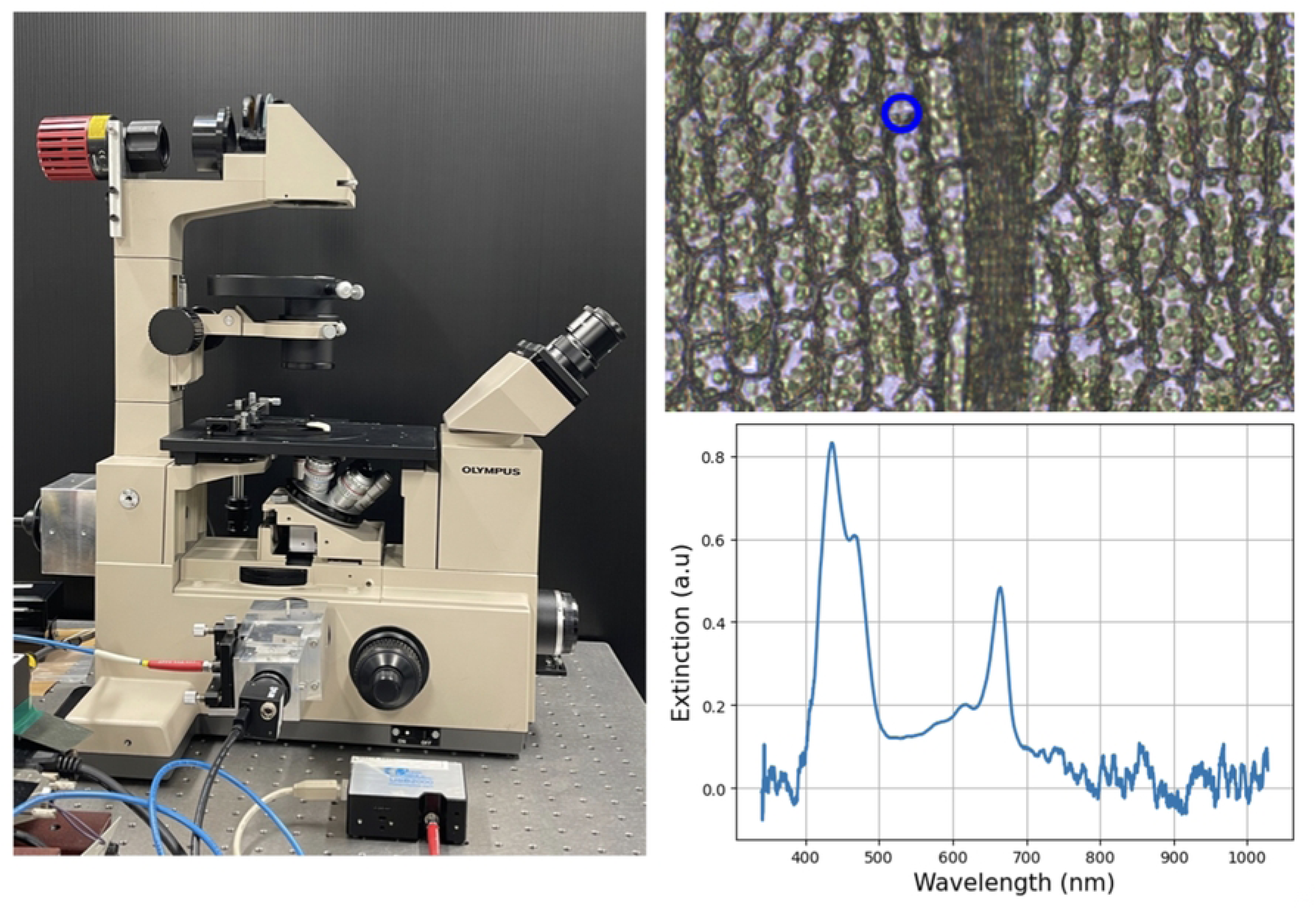
(Left) Photograph of the modified 2D+1D microscope; (Upper right) an example of the transmission microscopic image of moss leaf cell; (Lower right) the corresponding extinction spectrum of the region of interest (blue circle) within the sample.

**Figure 3.**
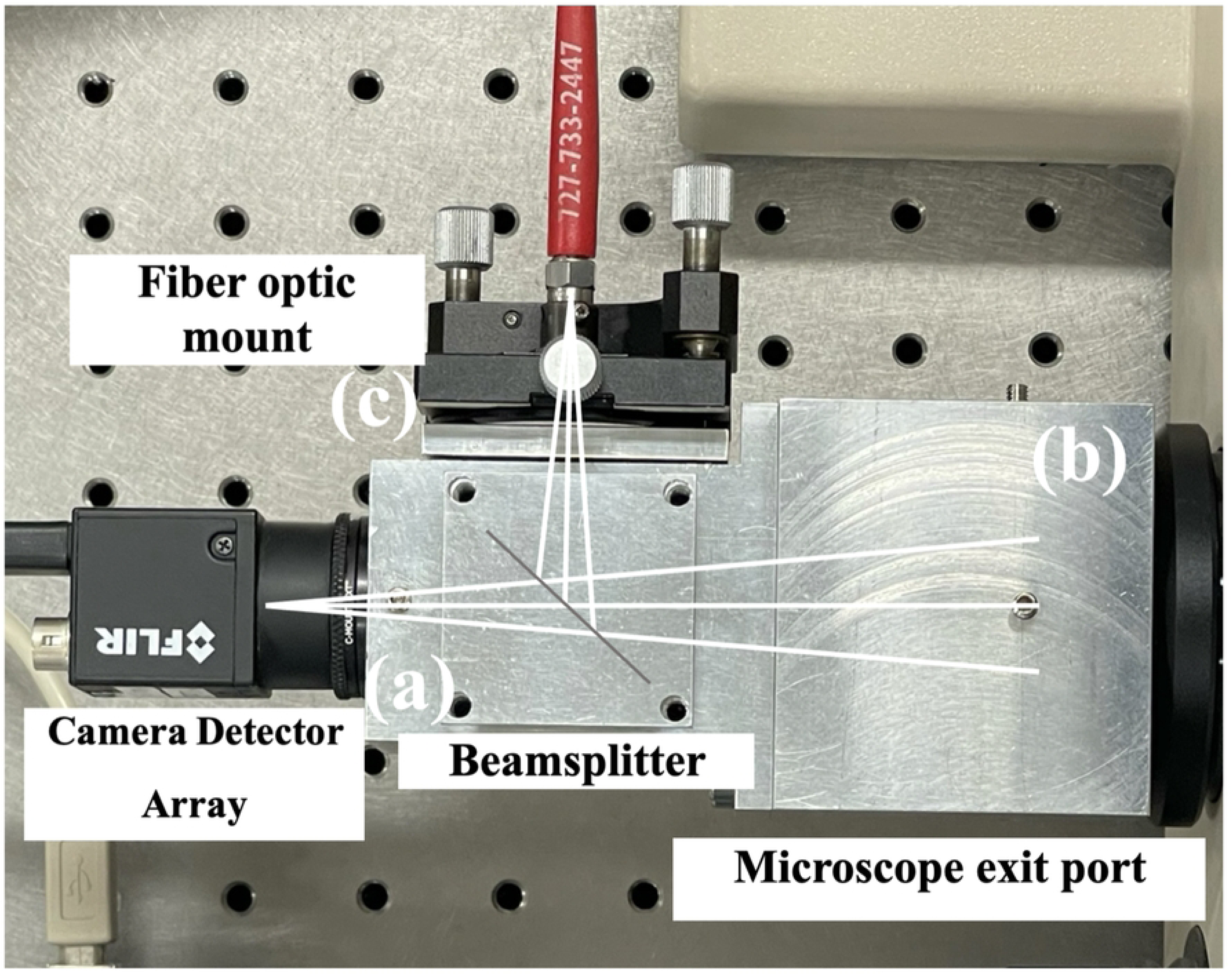
shows a close-up of the assembled 2D+1D adapter attached to the microscope exit port, along with an overlay of the optical rays aimed at the image center. The fully assembled system ensures the adapter is robust & portable, and allows easy adjustment of camera focus and fiber positioning during experiments.

**Figure 3a.** Close-up photograph of the 2D+1D adapter.

The optical components in our adapter consist of a 50/50 beam splitter, a fiber-coupled spectrometer (Ocean Optics USB2000, 350–1000nm spectral range, 1.4 nm resolution), and an RGB CMOS sensor (FLIR BFLY-U3-23S6C-C, 1920×1200 image, 5.86 µm pixels, 41 Hz).

For interactive use of the microscope system, we have developed software using Scientific Python that integrates the 2D camera image with the 1D spectral data in real time, while highlighting the spectrometer’s measurement region to the user (see Fig. 4). While the software allows saving 2D+1D video, the camera image and spectrometer data are captured asynchronously, so that there is some timing jitter to the acquisition, at the level of up to half the frame time. For samples that are moving slowly on the time scale of this frame time, this jitter has no apparent effect on the video data.

**Figure 4.**
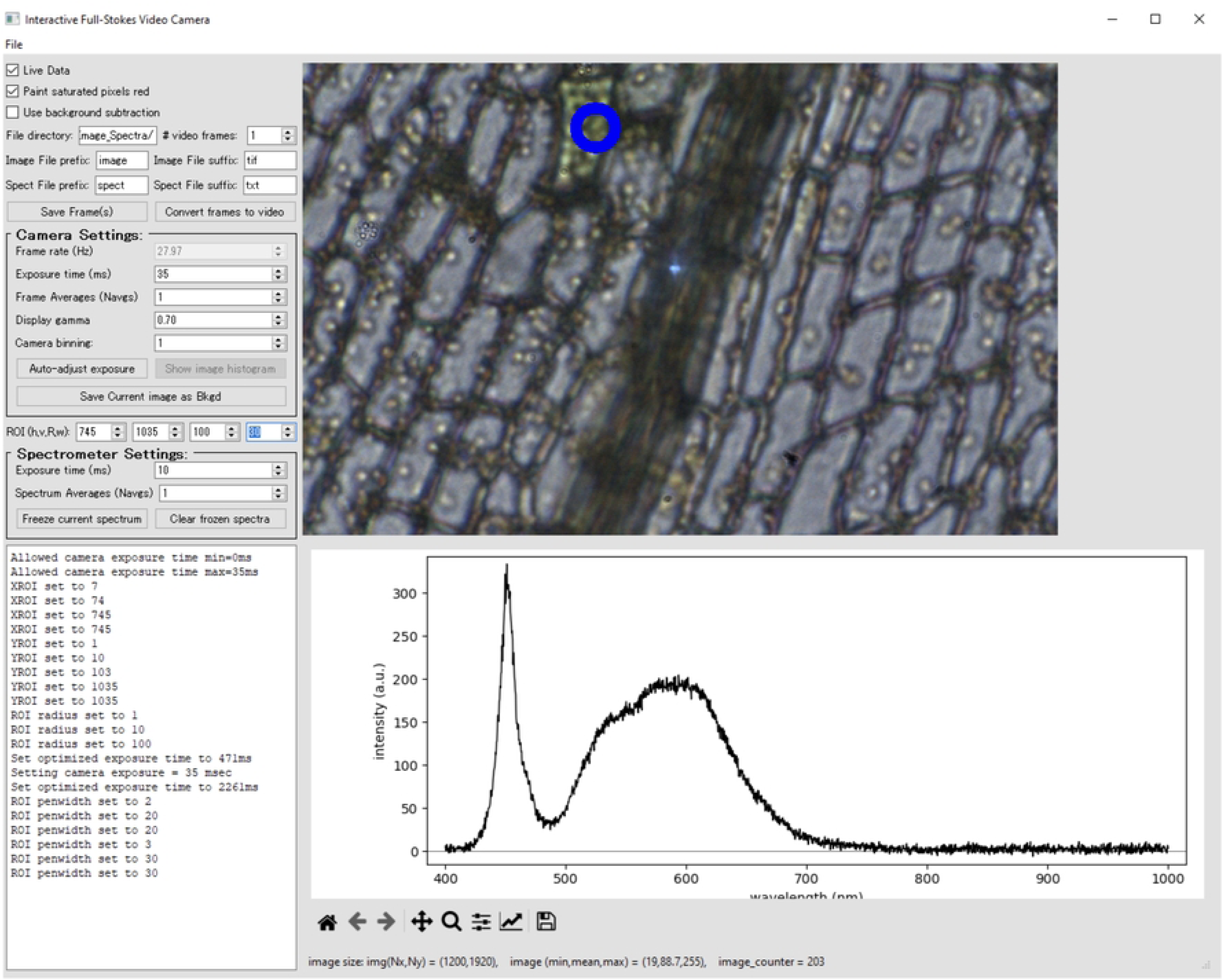
Our Python software interface, showing the 2D image and the corresponding spectral data measurement region indicated by a blue circle.

## 3. Measurement results and discussions

Based on the fully assembled system and the developed software, we investigate the 2D+1D imaging and spectral transmission and reflection characteristics of living cell samples, as a preliminary step toward applying other microscopy techniques (e.g., fluorescence imaging) in the future. The samples were prepared on glass slides and examined under a microscope with a 20X objective lens (Olympus LWD CDPlan 20x PL /0.40). By integrating spatial imaging with spectral information, structural features, chemical composition, and cell thickness variations can be revealed. The results demonstrate measurable thickness variations across multiple regions of interest (ROIs) within each sample, providing complementary and detailed insights into cellular characteristics [18].

First of all, the partial resolution of the microscope was characterized by imaging a USAF resolution target (PhotomaskPortal RT39D006), as shown in Fig. 5. Before adding the 2D+1D adapter, we measured the image resolution of the original microscope setup [Fig. 5(a)] with the 20X objective lens in transmission mode with white light LED illumination (MCWHLP1, 6500 K, 2350 mW (Min) Mounted LED, 700 mA), and in reflection mode with Xenon lamp illumination (Xenon lamp SLS605 (380nm - 780 nm)). We analyze the resolution target bars of element 6 in group 8 because it offers a suitable balance between small feature size and sufficient visibility. In group 9, the spacing between the narrower line pairs reduces intensity modulation making consistent contrast estimation difficult. The contrast of the resolution bars was calculated as *C* = (*I*_1_-*I*_0_)/(*I*_1_+*I*_0_) [19], where the maximum and minimum intensities *I*_1_ and *I*_0_ are taken from the peak-to-valley values within the image. Finally, we replaced the microscope’s original MTV-3 C-mount adapter with our 2D+1D adapter and took the same set of measurements in transmission and reflection [Fig. 5(b, c)]. The contrast measurements shown in Fig. 5 indicate that the two optical setups provide similar contrast values: 0.646 for the original microscope, and 0.716 for the 2D+1D setup. Unexpectedly, the 2D+1D results are better than the original microscope setup, but the difference of 0.070 is within the expected range of measurement variability for this contrast metric and the difficulty of accurately performing the contrast measurement. In terms of overall resolution, we have not been able to detect any significant differences between the two. This indicates that both setups provided comparable capability in distinguishing the dark and bright regions of the resolution target. For the contrast analysis, group 8, element 6, was selected because it offered a suitable balance between small feature size and sufficient visibility. In smaller groups or elements, the spacing between the line pairs became narrower, making the intensity modulation less distinct and the contrast separation more difficult. Therefore, group 8, element 6, was used as the smallest clearly resolvable feature for a reliable comparison between the two setups.

**Figure 5.**
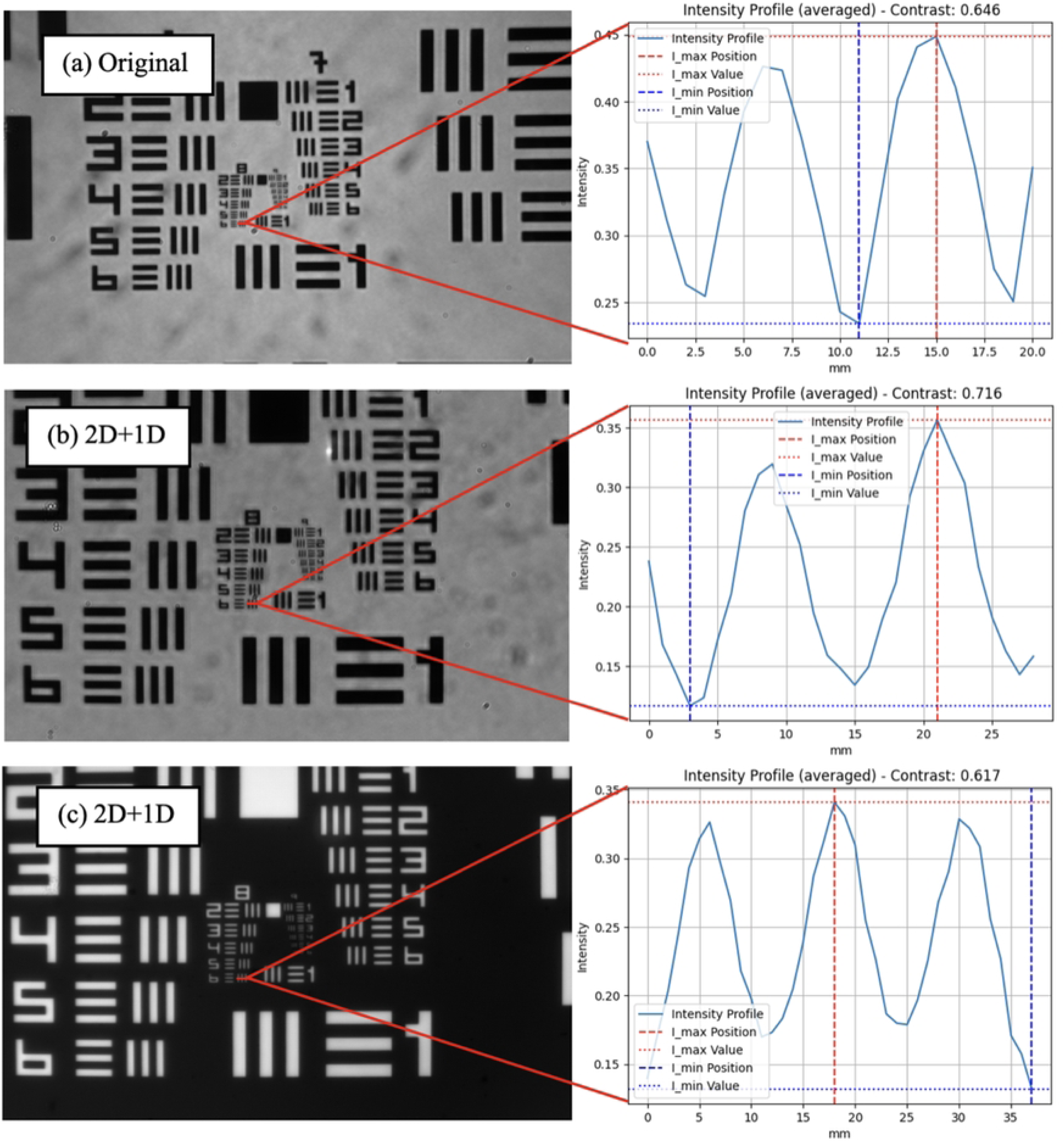
Optical performance characterization of the microscope with a 20X objective lens in (a) Image resolution of the original microscope setup, (b) transmission mode, and (c) reflection mode. No obvious differences in contrast or resolution were observed between the original microscope setup and the 2D+1D setup.

We used our integrated 2D+1D system to investigate living cells of the moss plant *Physcomitreium patens* and estimate chlorophyll concentration using spatial imaging and high-resolution spectral analysis [20, 21]. For chlorophyll estimation, calibration was required to account for factors such as optical path length and scattering effects in the measurement setup and sample. This calibration was performed using chlorophyll solutions with known standard concentrations. Chlorophyll was extracted from 10 mg of *Physcomitrium patens* gametophores using approximately 100 mL of 95% ethanol as the solvent to prepare a stock chlorophyll solution. After obtaining the solution, its concentration in mg/L was determined using a reference UV-Vis spectrometer (Ocean Optics USB4000, 350–1045 nm spectral range, 1.01 nm resolution) and calculated using equations which is used when chlorophyll is extracted with ethanol [22]

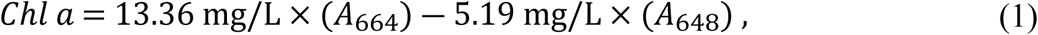

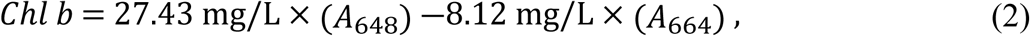

where *Chl a* represents chlorophyll a, *Chl b* represents chlorophyll b, *A*_664_ is the absorbance at 664 nm wavelength, and *A*_648_ is the absorbance at 648 nm. The total chlorophyll concentration is obtained by summing the values calculated from Eqs (1) and (2).

Measurements were performed in transmission mode, giving a sample transmission spectrum *I*(*λ*) (for the stock chlorophyll solution) and a reference transmission spectrum *I*_0_(*λ*) acquired under identical optical conditions using the solvent (80% ethanol) as the blank reference. The ratio *I*(*λ*)/*I*_0_(*λ*) represents the sample’s transmittance at each wavelength, corrected for the spectral profile of the light source and optical components. The absorbance spectrum is then calculated as [23]

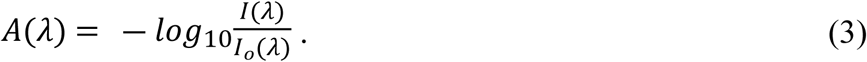

Chlorophyll absorbances *A*_664_ and *A*_648_ were evaluated by averaging the spectral data in the 663– 665 nm and 645–648 nm ranges, following established spectrophotometric chlorophyll determination methods [24]. The absorbance data of the stock chlorophyll solution (Fig. 6) were used to calculate the chlorophyll concentrations using Eqs (1) and (2), yielding chlorophyll a and chlorophyll b concentrations of 6.93 mg/L and 6.02 mg/L, respectively, for a total chlorophyll concentration of 12.95 mg/L.

**Figure 6.**
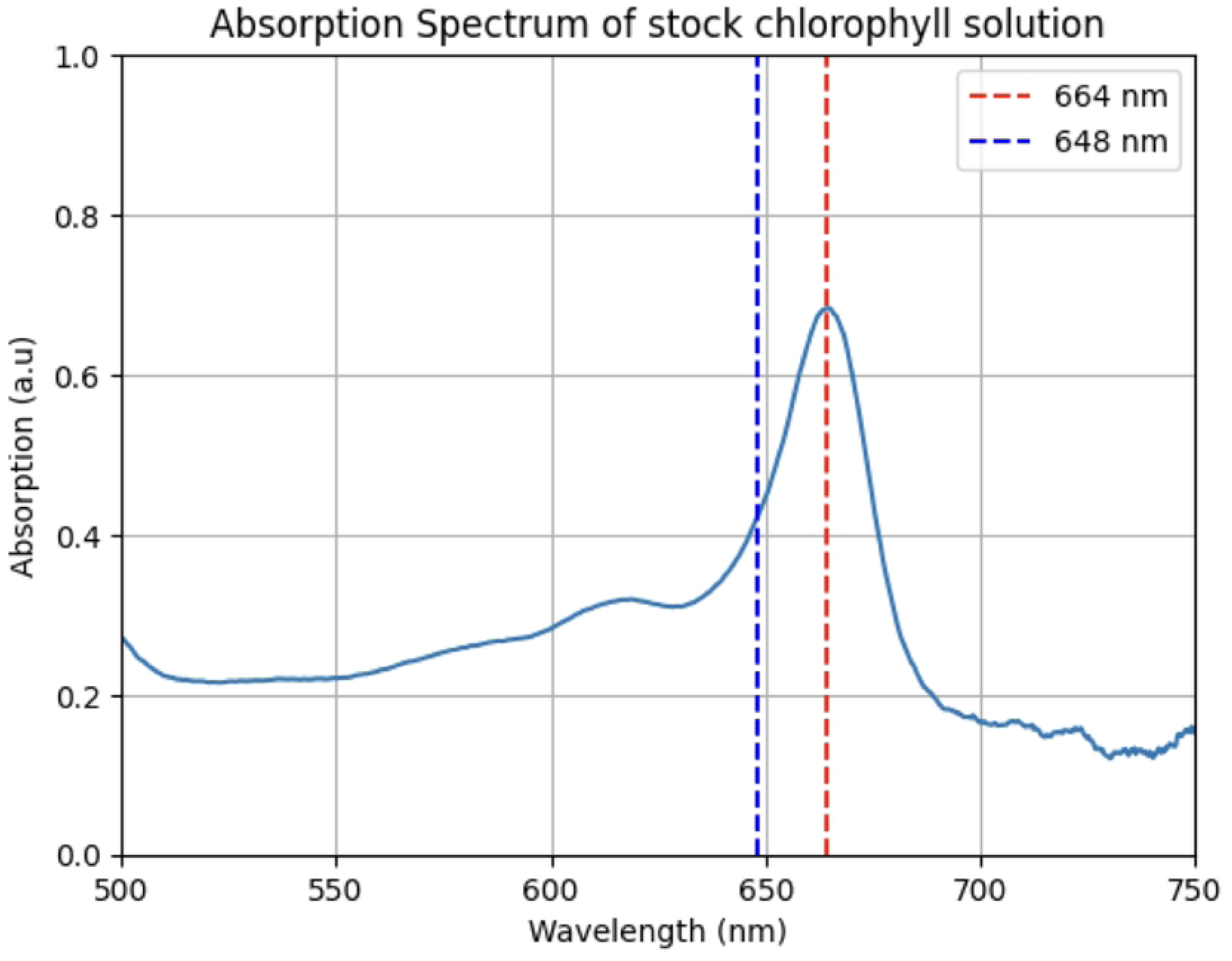
Absorbance spectrum of the stock chlorophyll solution.

The stock chlorophyll solution was also diluted to known standard concentrations for calibration of the 2D+1D spectral data. Standard solutions of 1, 2, 3, 4, and 5 mg/L were prepared using the dilution equation. *C*_1_*V*_1_ = *C*_2_*V*_2_for known concentrations *C* and mixing volumes *V*. Each standard solution was measured using the 2D+1D spectrometer under identical optical conditions (Fig. 8), from which we calculate the absorbance with Eq. 3. The known standard chlorophyll solutions (Fig. 7) were used as reference points to plot the calibration curve (Fig. 9), and we performed a linear regression between these known values and the measured absorption using the 2D+1D system spectrometer chlorophyll concentration measurements. This calibration curve shows a linear relationship between the spectrometer-measured absorption and the known chlorophyll concentration of the standard solutions. The regression equation is

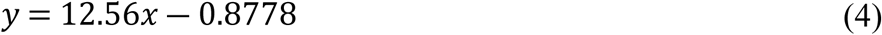

**Figure 7.**
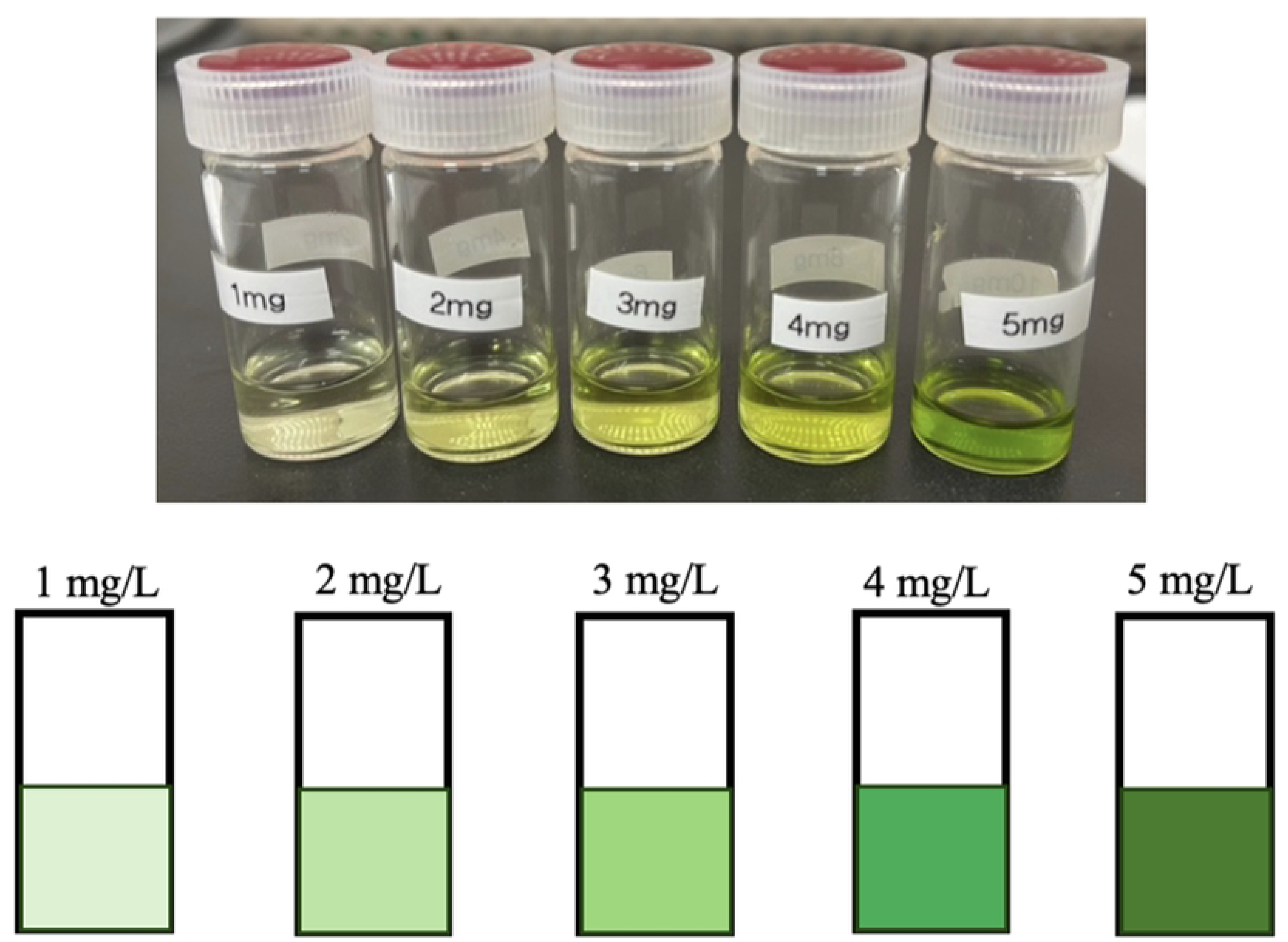
Standard of chlorophyll solution extracted from *Physcomitrium patens* moss for calibration curve used to fit a linear regression model.

**Figure 8.**
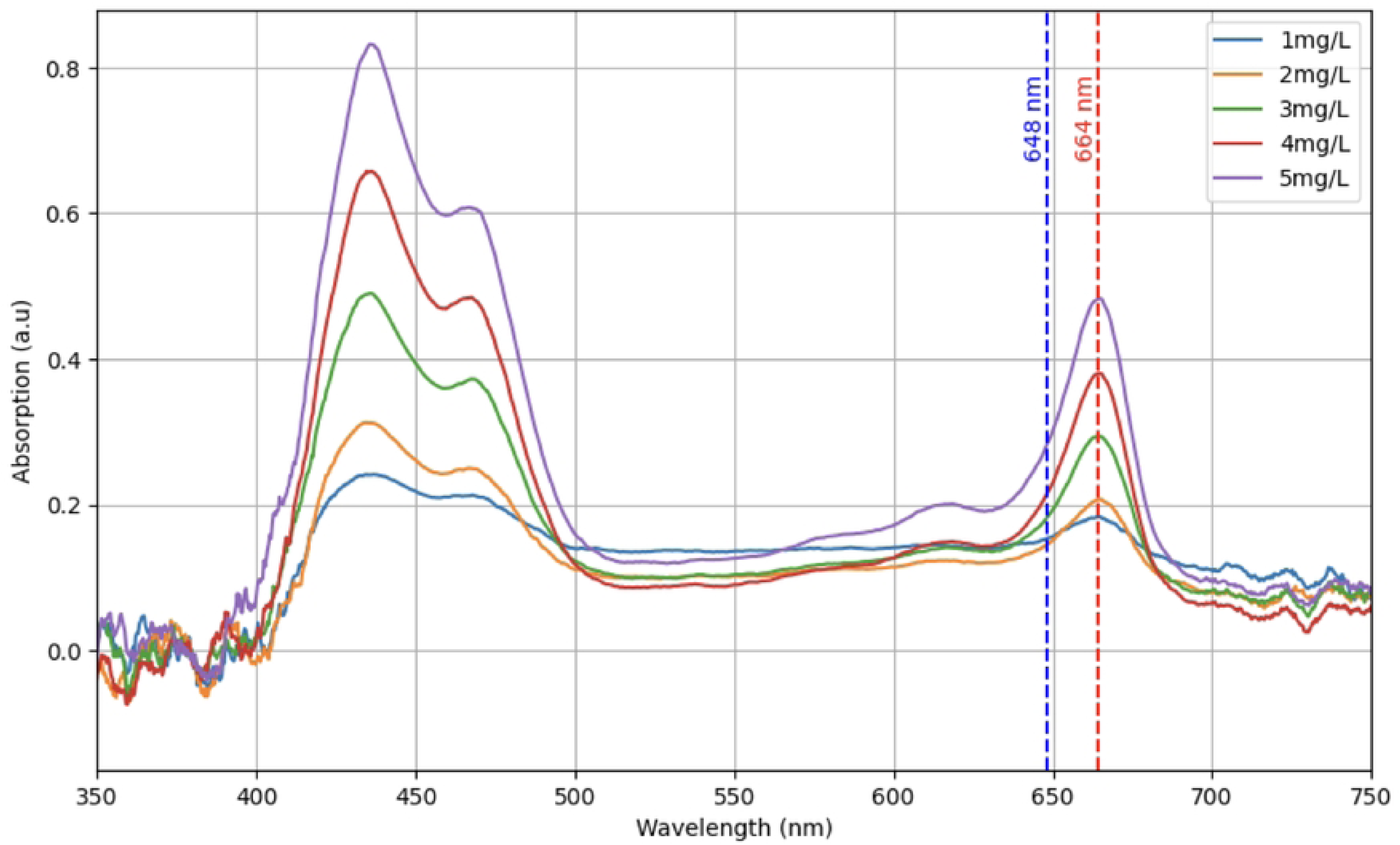
The absorbance spectra of the prepared standard chlorophyll solutions were measured by the 2D+1D spectrometer.

**Figure 9.**
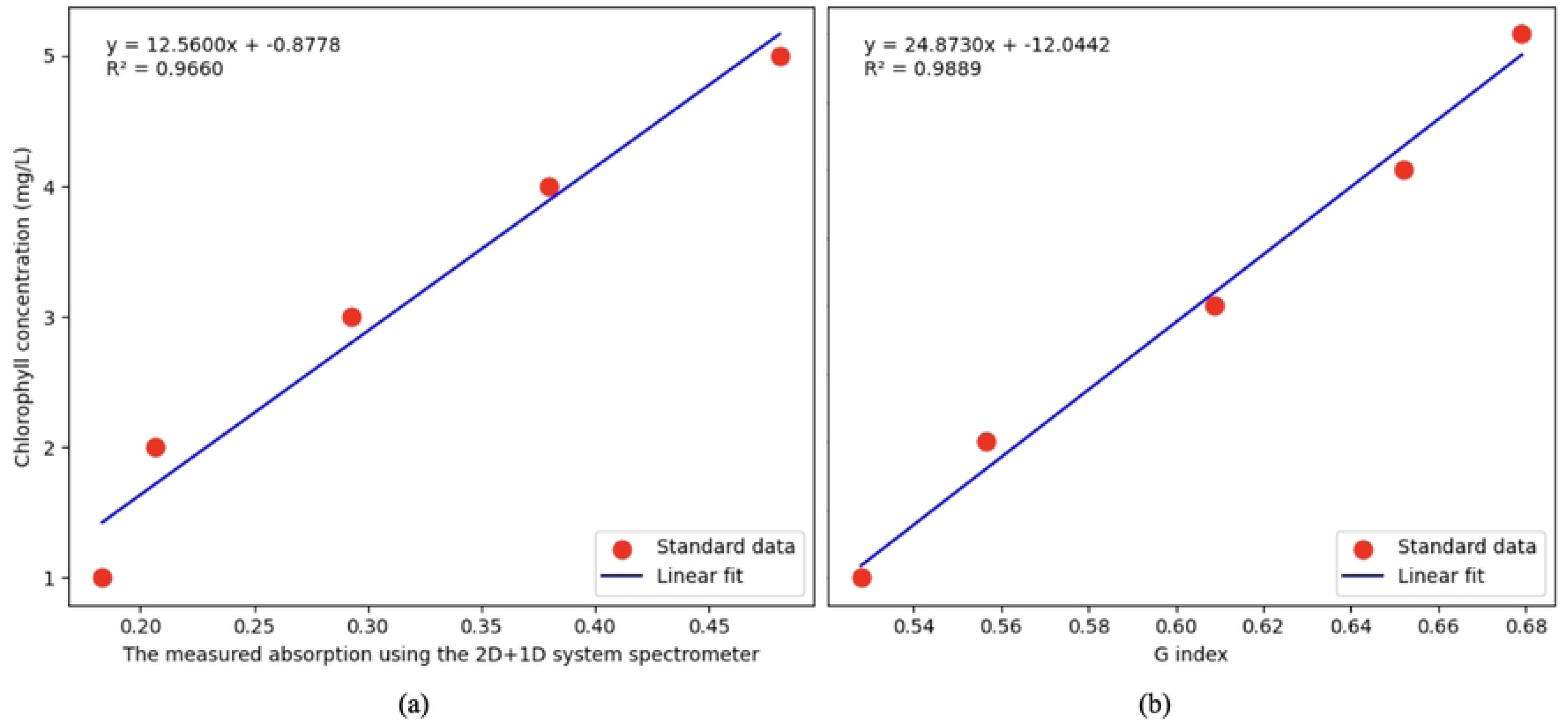
(a) The spectrometer calibration curve was obtained by plotting the known chlorophyll concentrations of standard solutions against the measured absorption spectra, (b) The color calibration curve obtained by plotting the known chlorophyll concentration of standard solutions with RGB index G/(R+B) from the 2D+1D system camera.

where *y* represents the chlorophyll concentration in mg/L and *x* represents the measured absorption using the 2D+1D system spectrometer. The coefficient of determination (*R)*^2^ was 0.9660, indicating approximate linearity between the measured absorbance and chlorophyll concentration [Fig.9(a)], as expected for thinly absorbing samples. With this calibration curve, we can convert measured absorbance of unknown samples to chlorophyll concentration in mg/L.

It is also possible to estimate the chlorophyll concentration from the image using the RGB index G/(R+B) at each pixel [25]. Using the same standard solutions, we generated a second calibration curve, for which the coefficient of determination (*R)*^2^ was 0.9892, as shown in Fig. 9(b). The RGB index regression equation is

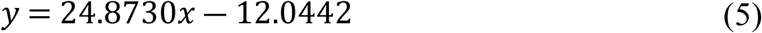

where *y* represents the estimated chlorophyll concentration in mg/L and *x* represents the RGB index G/(R+B).

Next, we observed the living leaf cells of *Physcomitrium patens* with the 2D+1D system. For sample preparation, we followed the previous paper [26]. In short, a 50-µm-thick silicone sheet was placed on a glass slide, and a 2–3 mm square region was excised from its center to form a 50-µm-deep chamber. Then, the chamber was filled with pure water. The gametophore leaf of Physcomitrium patens was excised under stereomicroscope, and the excised leaf was placed inside the chamber and covered with a coverslip. Example absorption spectra and transmittance images from the 2D+1D system are shown in Fig. 10. The corresponding measured absorption using the 2D+1D and RGB indices obtained from image and spectral data are presented in Table 1, together with the chlorophyll estimation results from both spectral analysis and RGB index, where the RGB index value is an average over the pixels within the corresponding ROI circle.

**Figure 10.**
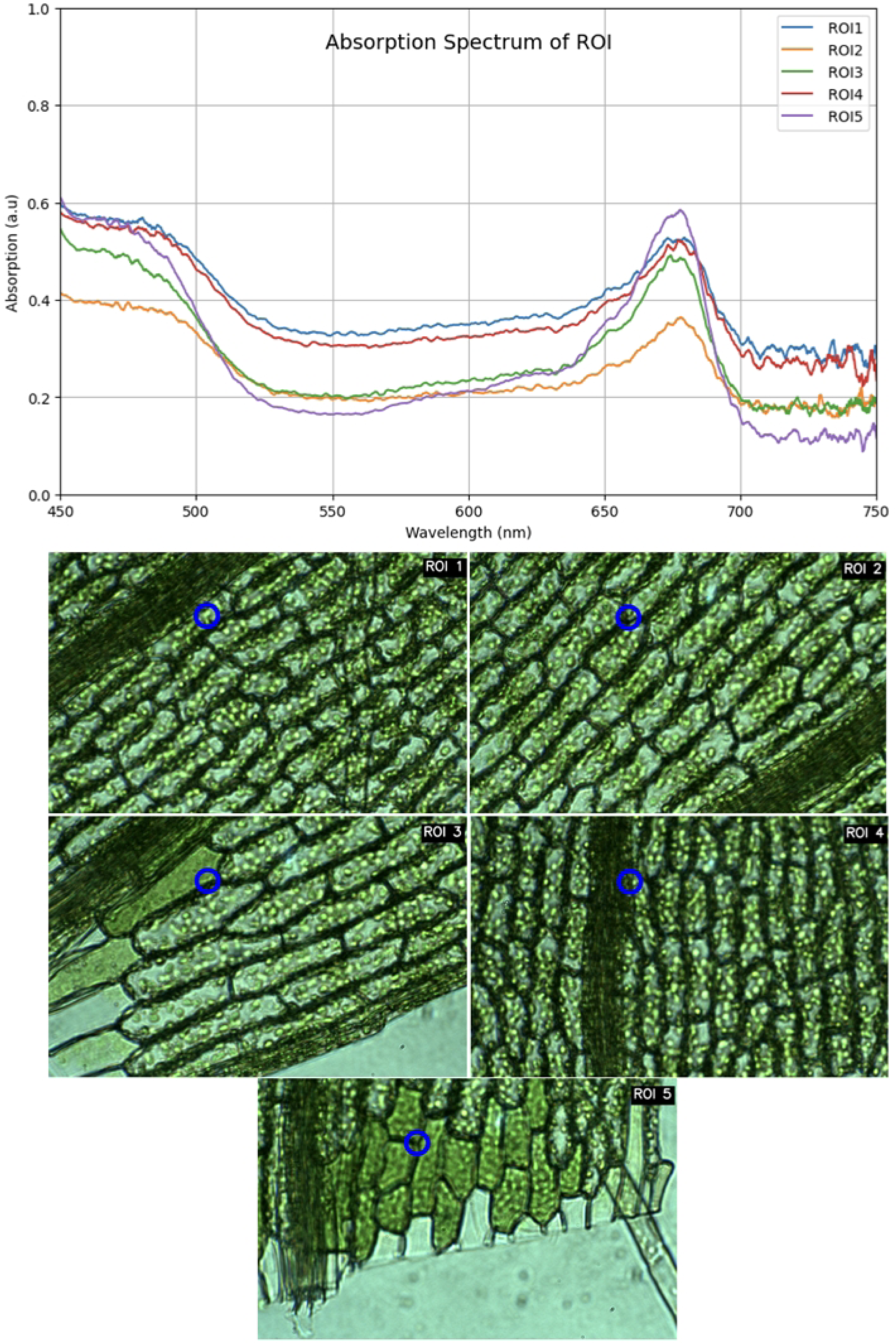
Example absorbance spectra and images of the leaf cells of *Physcomitrium patens* moss for estimating chlorophyll concentration. The spectral measurement regions for each image are shown as blue circles.

**Table 1.** Comparison of chlorophyll concentration estimates obtained using two methods.

| ROI | Chlorophyll concentration by Spectral | Chlorophyll concentration by Image |
| --- | --- | --- |
|  | Analysis (mg/L) | Analysis (mg/L) |
| 1 | 5.58 | 5.71 |
| 2 | 3.42 | 5.37 |
| 3 | 5.32 | 7.07 |
| 4 | 5.63 | 6.11 |
| 5 | 6.48 | 7.59 |

In Table 1, we see that the chlorophyll concentrations estimated by spectral and image analysis differ. Access to the spectrum is an advantage of our 2D+1D modification, and we believe this provides a more accurate estimate of chlorophyll because spectral analysis uses direct measurement, whereas image color analysis is indirect.

The radar plot (Fig. 11) compares the distribution patterns of total chlorophyll concentration obtained from spectrum-based and RGB image-based methods across five moss ROIs. All axes represent chlorophyll concentration (mg/L) on a scale of 0–9. Both methods showed a comparable overall pattern among the samples, indicating that RGB image analysis can reflect relative chlorophyll variation across the tested moss samples. The close geometric correspondence between the two radar plots, including similar shapes and relative proportions without crossover, suggests that both methods identify the same regions of higher and lower chlorophyll concentration. This demonstrates agreement between the two techniques in detecting spatial heterogeneity. However, the RGB image-based method produced consistently higher values than the spectrum-based method across all five ROIs, resulting in a uniformly larger polygon and indicating a systematic positive bias. The most pronounced relative discrepancy was observed in ROI 2, where the spectrum-based value reached its lowest concentration at 3.42 mg/L, compared with 5.37 mg/L from the RGB image-based method. In contrast, ROI 1 showed the smallest separation between methods, with a difference of 0.13 mg/L. Although minor differences in magnitude were observed, these variations did not alter the overall distribution pattern. Overall, the RGB-derived values followed the same general trend as the spectrum-derived values, supporting the use of RGB image analysis as a practical approach for reflecting relative differences in chlorophyll content among moss samples.

**Figure 11.**
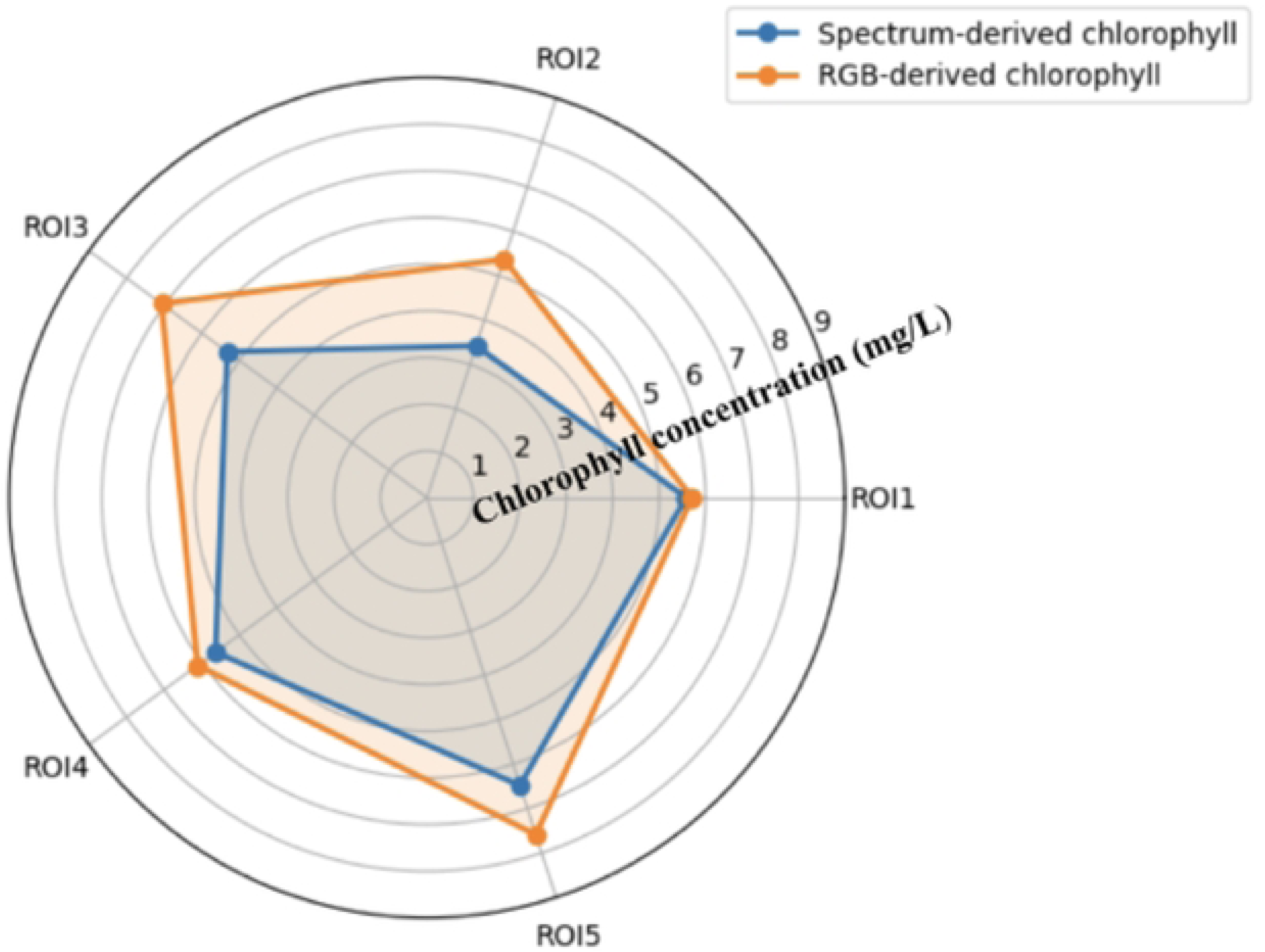
All axes: chlorophyll concentration (mg/L), scale 0-9. Image analysis yields a uniformly larger polygon than spectral analysis across all five ROIs, confirming a systematic positive bias ROI 2 shown the most pronounced relative discrepancy (spectral nadir at 3.42 mg/L vs image 5.37 mg/L).

In this study, we report chlorophyll concentration in mg/L, whereas in conventional plant physiology studies, chlorophyll content is often normalized to plant biomass and expressed as mg/g fresh weight or mg/g dry weight. Conversion between mg/L and mg/g requires the chlorophyll concentration in the extract, the final extraction volume, and the mass of plant material used. The Arnon-based calculation of total chlorophyll, for example, includes absorbance at 663–665 nm and 645–648 nm ranges, solvent volume, and sample fresh weight to express total chlorophyll as mg/g fresh weight [27]. The conversion from extract concentration *C* to biomass-normalized chlorophyll content can be expressed as: *C* (mg/g) = *C* (mg/L) × *V*/*m*, where *V* is the final extraction solvent volume, and *m* is the sample mass.

In fresh moss samples measured directly on glass slides, the chlorophyll-related spectral feature was observed in the red region, with a prominent signal near 680 nm. This spectral position differs from the absorption maxima typically used for solvent-extracted chlorophyll, where chlorophyll a and b are commonly evaluated around 664–665 nm and 648–649 nm in ethanol extracts. The shift toward the longer wavelength region may be associated with the in vivo pigment environment of chlorophyll within chloroplasts, tissue scattering, re-absorption, and possible chlorophyll fluorescence contribution. The red-region spectral response observed near 680 nm in fresh moss tissue is consistent with in vivo chlorophyll-related optical behavior rather than isolated chlorophyll in solvent. In intact photosynthetic tissues, chlorophyll molecules are bound to pigment–protein complexes within photosystems, and their spectral properties can differ from those of extracted pigments. In addition, chlorophyll fluorescence from photosystem II is commonly observed around 680–685 nm, while far-red fluorescence can appear near 730–740 nm. Therefore, the feature observed near 680 nm in fresh moss samples may reflect a combined contribution of chlorophyll absorption, tissue optical effects, re-absorption, and chlorophyll fluorescence [28]. This supports the need to calibrate the microscope-based measurement system using standards or reference spectra rather than relying solely on peak positions from solvent-extracted chlorophyll.

## 4. Conclusion

The 2D+1D adapter is a simple modification that attaches to a conventional microscope, allowing the simultaneous acquisition of wide-field imaging data together with spectral data from a single region of interest. The combination is simple and accessible, allowing users without specialized expertise to operate and implement the system. Although full hyperspectral imaging provides a more complete dataset of the sample, the 2D+1D approach offer simpler hardware and faster frame rates while still delivering the full spatial context of an image. For any location where the biological researcher wants to measure the spectrum, the user can move the sample to the desired location, and access much higher spectral resolution information than hyperspectral imaging instruments typically deliver. Ultimately, the proposed 2D+1D imaging and spectral approach enables interactive use during experiments, such that specific features within the sample can be tracked in real time. Having the microscope image provides the user insight into the spatial context of the measurement (e.g. how chlorophyll concentration varies across the cell and across the tissue) while spectrum provides a deeper insight into the chemistry of the sample (e.g. how much of the attenuation is due to chlorophyll and how much is due to scattering) that the sample’s color alone provides.

## Acknowledgements

This work was supported by the Japanese Government MEXT scholarship. This paper is an extended version of the conference proceedings paper [29].

## Supporting information

1. CAD files mechanical components of the adapter: (a) the beamsplitter mount, (b) the exit port coupler, and (c) the fiber mount coupler.

